# Hippocampus shape across the healthy lifespan and its relationship with cognition

**DOI:** 10.1101/2020.10.30.362921

**Authors:** Aurelie Bussy, Raihaan Patel, Eric Plitman, Stephanie Tullo, Alyssa Salaciak, Saashi A. Bedford, Sarah Farzin, Marie-Lise Béland, Vanessa Valiquette, Christina Kazazian, Christine L. Tardif, Gabriel A. Devenyi, Mallar Chakravarty

**Affiliations:** Computational Brain Anatomy (CoBrA) Laboratory, Cerebral Imaging Centre, Douglas Mental Health University Institute, Montreal, Quebec, Canada; Integrated Program in Neuroscience, McGill University, Montreal, Canada; Department of Psychiatry, McGill University, Montreal, Quebec, Canada; Department of Biomedical Engineering, McGill University, Montreal, Quebec, Canada; McConnell Brain Imaging Centre, Montreal Neurological Institute, Montreal, Quebec, Canada; Department of Neurology and Neurosurgery, McGill University, Montreal, Quebec, Canada

## Abstract

The study of the hippocampus across the healthy adult lifespan has rendered inconsistent findings. While volumetric measurements have often been a popular technique for analysis, more advanced morphometric techniques have demonstrated compelling results that highlight the importance and improved specificity of shape-based measures. Here, the MAGeT Brain algorithm was applied on 134 healthy individuals aged 18-81 years old to extract hippocampal subfield volumes and hippocampal shape measurements, notably: local surface area (SA) and displacement. We used linear, second or third order natural splines to examine the relationships between hippocampal measures and age. In addition, partial least squares analyses were performed to relate measurements with cognitive and demographic information. Volumetric results indicated a relative preservation of the right cornus ammonis 1 with age and a global volume reduction linked with older age, female sex, lower levels of education and cognitive performance. Vertex-wise analysis demonstrated an SA preservation in the anterior hippocampus with a peak during the sixth decade, while the posterior hippocampal SA gradually decreased across lifespan. Overall, SA decrease was linked to older age, female sex and, to a lesser extent lower levels of education and cognitive performance. Outward displacement in the lateral hippocampus and inward displacement in the medial hippocampus were enlarged with older age, lower levels of cognition and education, indicating an accentuation of the hippocampal “C” shape with age. Taken together, our findings suggest that vertex-wise analyses have higher spatial specifity and that sex, education and cognition are implicated in the differential impact of age on hippocampal subregions throughout its antero-posterior and medial-lateral axes.

## 1. Introduction

The role of the hippocampus in normal behaviour and across neuropsychiatric disorders has been a major topic of clinical and neuroscientific research since the initial investigations of Scoville and Milner on patient H.M. (Scoville and Milner 1957). Since these pioneering investigations, altered hippocampal structure and function have been implicated in several neuropsychiatric disorders, such as Alzheimer’s disease [AD, (Convit et al. 1997; Amaral et al. 2018; Sabuncu et al. 2011)], schizophrenia (Nelson et al.1997), major depressive disorder (J. D. Bremner et al. 2000; Videbech and Ravnkilde 2004; Treadway et al. 2015), posttraumatic stress disorder (J. Douglas Bremner et al. 2003; Apfel et al. 2011), and frontotemporal dementia (Laakso et al. 2000; Muñoz-Ruiz et al. 2012; Chapleau et al. 2020). Moreover, the relationship between hippocampal structure and age across the adult lifespan has also been of interest towards setting a “baseline” of hippocampal morphology, against which disorders implicating this region may be compared. However, previous findings investigating hippocampal changes with age have rendered inconsistent results (La Joie et al. 2010; de Flores, La Joie, and Chételat 2015; A. Bussy et al. 2020) and potential sources of this inconsistency are discussed below.

Although whole hippocampal volume can be studied using standard volumetric magnetic resonance imaging (MRI; (Sabuncu et al. 2011; Sankar et al. 2017; Schoemaker et al. 2016)), there is mounting evidence that variation at the level of the hippocampal subfields provides additional insight into neuropsychiatric disorders and healthy aging. However, it is presently unclear which subregions of the hippocampus are the most impacted through the course of the adult lifespan. Indeed, some studies have demonstrated a predominant decrease of the cornus ammonis (CA)1 and subiculum volumes with age (Shing et al. 2011; Wisse et al. 2014; de Flores et al. 2015; Wolf et al. 2015; Daugherty et al. 2016), while others have shown a relative preservation of the CA1 volume compared to global hippocampal atrophy with age (Amaral et al. 2018; A. Bussy et al. 2020). These contradictory results might potentially be explained by the use of various hippocampal subfield definitions and segmentation techniques across existing literature (Yushkevich et al. 2015; Wisse et al. 2017; A. Bussy et al. 2020).

To date, studies investigating the hippocampus across the adult lifespan have largely been limited to the examination of volumetric information. However, different morphological techniques have been shown to provide spatially localized information about age-related hippocampal modifications, independent of finer-grain neuroanatomical definitions (Yang et al. 2013; Voineskos et al. 2015). For example, some groups have reported inward deformations in the hippocampal head of older adults (Yang et al. 2013). Additionally, voxel-based regression analyses demonstrated that age-related variation occurred mostly in the head and tail of the hippocampus (Pruessner et al. 2001). Furthermore, significant patterns of inward displacement modifications in the hippocampal head and outward displacement in the hippocampal body have been found to be associated with age (Voineskos et al. 2015). In contrast, another study found a higher impact of age in the posterior hippocampus rather than in the hippocampal head (Kalpouzos et al. 2009).

There are several other factors that may impact findings from previous studies. For example, previous normative aging studies have examined various age ranges, with some researchers considering large spans of the adult lifespan (Mueller et al. 2007; de Flores et al. 2015; Amaral et al. 2018; F. Zheng et al. 2018; A. Bussy et al. 2020), while others only investigated elderly individuals (Frisoni et al. 2008; Wisse et al. 2014). Despite previous analyses showing non-linear relationships between brain volume and age (Coupé et al. 2017; Tullo et al. 2019; A. Bussy et al. 2020), non-linear relationships with age are still rarely investigated, especially in morphometric studies (Yang et al. 2013).

The role of the hippocampus in episodic and working memory has previously been established (Driscoll et al. 2003; Eichenbaum 2004). Additionally, the protective role of high level of education (Noble et al. 2012) and cognition (Vuoksimaa et al. 2013) on the hippocampus have been demonstrated. Researchers have also established the impact of sex on disorders where hippocampal dysfunction and degeneration are implicated (Fleisher et al. 2005; Abel, Drake, and Goldstein 2010; Fisher, Bennett, and Dong 2018). Moreover, apolipoprotein E4 (APOE4) genotype has been recognized as a genetic risk factor for hippocampal age-related and AD-related atrophy (Pievani et al. 2011; O’Dwyer et al. 2012). However, to our knowledge, all these factors have rarely been studied in relation to hippocampal shape (Voineskos et al. 2015).

To extend our understanding of hippocampal subfield volumetry and morphology throughout the lifespan, the aim of this paper is to investigate the associations between age and hippocampal structure. We further investigate the relationship with different factors known to influence age-related variation such as biological sex, education, and APOE4 genotype. Finally, we also use a multivariate technique to connect hippocampal morphometry, demographic characteristics, and cognition. To the best of our knowledge, this presents a novelty within the field given that previous work typically investigates hippocampal morphometry using univariate analyses, linear models, or group comparisons to principally analyse the effect of one variable at the time (Wang et al. 2003; Yang et al. 2013; Voineskos et al. 2015; Dong et al.2019). Here, we explore the interaction of different variables at the same time, which allows us to draw conclusions about the influence of each variable towards overall shape modification.

## 2. Methods

### 2.1. Participants

Participants were selected from two datasets collected by our group; namely, the Healthy Aging (HA) and Alzheimer’s Disease Biomarkers (ADB) datasets (as previously described in our study investigating the striatum, globus pallidus, and thalamus in (Tullo et al. 2019) and another examining the effects of MR sequence on hippocampal subfield volumes estimates (A. Bussy et al. 2020)). Both HA and ADB datasets were approved by the Research Ethics Board of the Douglas Research Centre, Montreal, Canada and written informed consent was collected from all participants. 112 healthy individuals aged 18 to 80 (55 males and 57 females) and 66 healthy seniors aged 56 to 81 (25 males and 41 females) were selected from the HA and ADB datasets, respectively. Exclusion criteria included having a history of neurological disease, psychiatric disorder, major structural neurological abnormalities or pathologies, history of brain damage or concussion, current or recent use of psychoactive substances, intellectual disability, a Mini Mental State Examination (MMSE; (Folstein, Folstein, and McHugh 1975)) score below 24, having less than 6 years of formal education, and any contraindications to MRI.

The initial dataset, which is described in Supplementary table 1, summarizes the demographic information of the 178 participants initially included in the study. Table 1 describes the demographic information of the 134 individuals (“Full” dataset) for which the preprocessing and processing steps passed our quality control (QC; see 2.3. Raw quality control and 2.7. Output quality control). These participants were included for the age analyses using vertex-wise linear mixed-effect analyses (See 2.9. Statistics). Table 1 also includes demographics for the 85 individuals (“APOE4” dataset) who passed QC and with known APOE4 genotype, repeatable battery for the assessment of neuropsychological status (RBANS; (Randolph et al. 1998)), MMSE, and years of education. These participants were included in subfield-wise hippocampal volume analyses and partial least square (PLS) analyses (See 2.9. Statistics).

**Table 1.** Complete demographic information of the 85 individuals who passed motion and segmentation QC and with complete APOE4 genotyping, RBANS total score, MMSE, and years of education by dataset and demographics of the “Full” dataset with the 134 participants used for the vertex-wise age analyses.

|  | APOE4 - 85 participants |  |  | FULL - 134 participants |  |  |
| --- | --- | --- | --- | --- | --- | --- |
|  | ADB<br>(n=44) | HA<br>(n=41) | Overall<br>(n=85) | ADB<br>(n=54) | HA<br>(n=80) | Overall<br>(n=134) |
| **Age** |  |  |  |  |  |  |
| Mean<br>(SD) | 70.0<br>(5.64) | 51.6<br>(17.4) | 61.1<br>(15.7) | 69.9<br>(5.4) | 47.2<br>(16.2) | 56.4<br>(17.1) |
| Median<br>[Min, Max] | 70.0<br>[57.0, 81.0] | 50.0<br>[18.0, 80.0] | 66.0<br>[18.0, 81.0] | 70.0<br>[57.0, 81.0] | 46.0<br>[18.0, 80.0] | 62.5<br>[18.0, 81.0] |
| **Sex** |  |  |  |  |  |  |
| Female | 28 (63.6%) | 23 (56.1%) | 51 (60.0%) | 32 (59.3%) | 46 (57.5%) | 78 (58.2%) |
| Male | 16 (36.4%) | 18 (43.9%) | 34 (40.0%) | 22 (40.7%) | 34 (42.5%) | 56 (41.8%) |
| **APOE4** |  |  |  |  |  |  |
| Non-carriers | 30 (68.2%) | 28 (68.3%) | 58 (68.2%) |  |  |  |
| Carriers | 14 (31.8%) | 13 (31.7%) | 27 (31.8%) |  |  |  |
| **Education** |  |  |  |  |  |  |
| Mean (SD) | 16.4 (3.98) | 16.7 (3.42) | 16.6 (3.70) |  |  |  |
| Median [Min, Max] | 16.0 [10.0, 31.0] | 17.0 [10.0, 26.0] | 16.0 [10.0, 31.0] |  |  |  |
| **RBANS** |  |  |  |  |  |  |
| Mean (SD) | 95.0 (12.7) | 103.0 (15.9) | 98.7 (14.8) |  |  |  |
| Median [Min, Max] | 92.5 [71.0, 123.0] | 104.0 [57.0, 132.0] | 99.0 [57.0, 132.0] |  |  |  |
| **MMSE** |  |  |  |  |  |  |
| Mean (SD) | 28.6 (1.53) | 28.7 (1.54) | 28.6 (1.53) |  |  |  |
| Median [Min, Max] | 29.0 [24.0, 30.0] | 29.0 [24.0, 30.0] | 29.0 [24.0, 30.0] |  |  |  |

### 2.2. Imaging acquisition

All participants were scanned on a 3T Siemens Tim Trio MRI scanner using a 32-channel head coil at the Douglas Research Centre, Montreal, Quebec, Canada. T1-weighted (T1w) images were acquired using parameters established by the Alzheimer’s Disease Neuroimaging Initiative (ADNI) magnetization prepared - rapid acquisition by gradient echo (MPRAGE) protocol (Jack et al. 2008). T1w acquisition parameters were TE/TR = 2.98 ms/2300 ms, TI = 900 ms, α = 9°, 256 × 240 × 176 matrix, GRAPPA of 2, bandwidth = 238 Hz/pixel, 1.00 mm isotropic voxel dimensions, and scan time 5:12.

### 2.3. Raw quality control

Involuntary movements such as cardiac or respiratory motion can lead to motion artifacts and may negatively impact the quality of structural MRI images (Bellon et al. 1986; Smith and Nayak 2010; Reuter et al. 2015) and any quantitative outputs derived from these images (Reuter et al. 2015; Alexander-Bloch et al. 2016). Quality control (QC) of all raw images was performed by one of the authors (AB) using the QC procedure previously developed in our laboratory (Bedford et al. 2020); https://github.com/CoBrALab/documentation/wiki/Motion-Quality-Control-Manual).

### 2.4. Preprocessing

Image preprocessing was used to standardize the overall quality of the T1w images prior to processing. The minc-bpipe-library pipeline (https://github.com/CobraLab/minc-bpipe-library) was applied to T1w images and consists of several steps, including: N4 bias field correction (Tustison et al. 2010), registration to Montreal Neurological Institute (MNI) space (ICBM 2009c Nonlinear Symmetric) using bestlinreg (Collins et al. 1994; Dadar et al. 2018), and standardization of the field-of-view by using a MNI head mask transformed into native space using the linear transformation estimated at the prior stage. Finally, any voxels outside the head mask were cropped, the orientation of the brain was standardized, and the brain was extracted using Brain Extraction based on nonlocal Segmentation Technique [BEaST (Eskildsen et al.2012)].

### 2.5. Volumetric analysis

Segmentation of the hippocampal subfields was performed using the Multiple Automatically Generated Templates (MAGeT) Brain algorithm (Pipitone et al. 2014; M. M. Chakravarty and Steadman 2013). Grey matter subfields included the CA1, combined CA2 and CA3 (CA2CA3), combined CA4 and dentate gyrus (CA4DG), stratum radiatum/lacunosum/moleculare (SRLM) and subiculum; these were defined in five high-resolution atlases. These atlases have been previously manually segmented on 0.3 mm isotropic T1w and T2w MRI-based atlases (Winterburn et al. 2013) and the segmentation procedure was validated using both manual segmentation, comparisons to other pipelines, and simulations (Pipitone et al. 2014).

As described in our recent work (A. Bussy et al. 2020), MAGeT Brain was used to segment each dataset independently, with a first run for ADB dataset and another for HA dataset. First, a “best template selection” stage (https://github.com/CoBrALab/documentation/wiki/Best-Templates-for-MAGeT) was performed in order to select the 21 subjects with the highest quality of atlas-to-template segmentation in each dataset independently. Next, these 21 subjects were used to populate the template library in order to segment all the subjects of each dataset. This step improves the quality of the atlas-to-template segmentation by artificially increasing the number of candidate labels to 105 (21 templates x 5 atlases). A majority vote technique was then used on these 105 candidate labels to obtain final labels (Pipitone et al. 2014; M. M.Chakravarty and Steadman 2013; Makowski et al. 2018), which, when coupled with the atlas inflation strategy, performs similarly to or exceeds performance of recent advances in label fusion techniques (Bhagwat et al. 2016). Affine and SyN nonlinear registrations from the Advanced Normalization Tools (ANTS; [(Avants et al. 2008)]) are used within the MAGeT brain algorithm. Regions-of-interest located at each hippocampal subfield label were used to create single value masks. These masks were then dilated with a 3 mm radius kernel in order to focus both affine and nonlinear registration, a method which reduces computational time and improves segmentation accuracy (M. Mallar Chakravarty et al. 2008, 2009).

### 2.6. Shape analysis

The MAGeT pipeline also provides morphometric (surface area [SA] and displacement) measurements (Raznahan et al. 2014; Voineskos et al. 2015; Shaw et al. 2014; Janes et al. 2015). An average neuroanatomical representation was created for the hippocampal GM through group-wise nonlinear registration of the five atlases previously defined in (Winterburn et al. 2013); this representation was referred to as the “model” (Figure 1A). This model provides a common space for analysis of surface-based metrics from which we derived a mesh of the hippocampal surfaces, composed of ~1200 vertices/hemisphere. Then, a surface-based metric (Lerch et al. 2008) was used to estimate the local shape variation of our subjects with the model as a reference. Finally, SA (examining expansion or contraction) and displacement (examining inward or outward) were calculated using vertex-wise analysis (Figure 1B and 1C).

**Figure 1:**
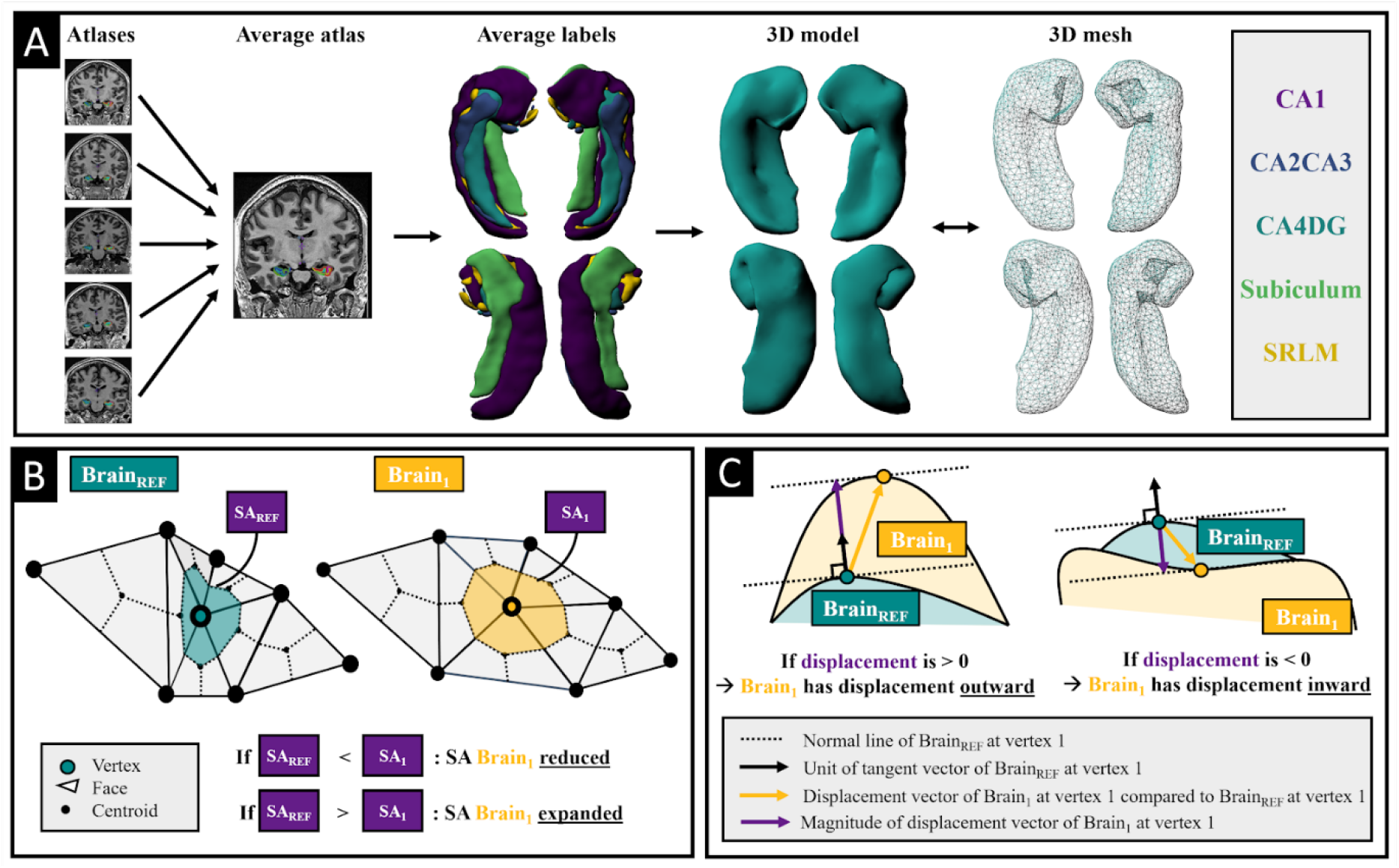
Schematic representation of the major analysis streams presented. **A)** The five hippocampal atlases were non-linearly registered to create an average brain. The same transformations were applied on the five hippocampal labels to obtain the five atlases in a common space, and labels were fused in common space using majority vote. An average 3D hippocampal model was created along with a 3D mesh in order to perform surface-based morphological analyses. **B)** Surface area (SA) was calculated using the Voronoï method of parcellation around each vertex. The SA of the brain of interest was then compared to the SA of the corresponding vertex in the reference brain. **C)** Displacement measure was calculated using the magnitude of the displacement vector between a vertex and the corresponding vertex in the reference brain.

### 2.7. Output quality control

QC of the final labels by visual inspection was conducted following the procedure implemented by our group (https://github.com/CobraLab/documentation/wiki/MAGeT-Brain-Quality-Control-(QC)-Guide). All QC was conducted by one of the authors (AB) to only include high quality segmentation in the statistical analyses.

### 2.8. Subfields objects

From the five brains used previously to define our five atlases (Winterburn et al. 2013), average brains and labels were previously created (Voineskos et al. 2015). Mango software (*version 4.1 developed by Research Imaging Institute, UTHSCSA*; (Lancaster et al. 2011)) was used to create 3D objects of each subfield to visually help comparison between our morphological measurements and the hippocampal subfields (Figure 1A).

### 2.9. Statistics

#### 2.9.1. Age-relationship models

First we sought to examine the impact of age on our morphometric and volumetric measures while accounting for factors like sex, genetic risk, education and cognition. The “APOE4” dataset of 85 individuals (Table 1) was used to investigate the hippocampal subfield volume changes with age while the “FULL” dataset of 134 individuals was used to examine the relationship between the vertex-wise morphological measures and age. This allowed us to take advantage of the higher number of participants to extract vertex-wise relationships with age. Linear mixed-effects models (lmer from *lmer Test_3.1-2* package for subfield-wise analysis or *vertexLmer* from *RMINC_1.5.2.2* package for vertex-wise analysis in R 3.6.3) and natural splines (ns from *splines* package) were used to model linear, second, or third order age relationships, either for subfield-wise volumes or vertex-wise measurements. Akaike information criterion [AIC; (Akaike 1974)] was used to investigate the most appropriate age relationships across the multiple models. The model with the lowest AIC was selected and was considered to best fit the data (Mazerolle 2006); see also previous work from our group (Tullo et al. 2019; Bedford et al. 2020). All statistical models are described in detail within the supplementary material.

For the subfield-wise analysis, sex, intracranial volume, APOE4 status, education, MMSE, and RBANS were included as fixed effects, while dataset was included as a random effect. Bonferroni correction was performed to correct for multiple comparisons across our 12 structures (bilateral total hippocampus and five GM subfields per hemisphere), at a p<0.05 threshold for significance, resulting in a significance level of p<0.00417 (only corrected p-values reported throughout this paper).

For the vertex-wise analysis, sex was included as a fixed effect, while dataset was included as a random effect. A 5% false-discovery rate (FDR) correction was applied to control for the expected proportion of “discoveries” that are false (Yoav Benjamini and Hochberg 1995; Y. Benjamini et al. 2001). After selecting the best fit for each vertex, linear mixed-effects models (lmer from *lmerTest_3.1-2* package in R 3.6.3) and *which.max* and *which.min* functions were used to calculate the age for which SA and displacement were at the maximum and minimum, respectively.

#### 2.9.2. Partial least squares analysis

After establishing the baseline with more common univariate measures, we sought to examine how morphometry and volume covaries with factors that we know are implicated in the ageing process, such as sex, genetic risk, education and cognition. Partial least squares (PLS), a multivariate technique that transforms the predictors to a smaller set of uncorrelated components and subsequently performs least squares regression on these newly-defined components, was performed on the APOE4 dataset. The goal of PLS is to identify a set of latent variables (LVs), that explain patterns of covariance between “brain” and “cognitive/demographic” data with the constraint that LVs explain as much of the covariance between the two matrices as possible. Mathematically, each LV will describe linear combinations of the “brain” and “cognitive/demographic” data that maximally covary. Two separate PLS analyses were used; the first to investigate the relationships between the hippocampal subfields volumes and cognitive/demographic information, and the second to relate shape measurements with cognitive/demographic information. In this work, we first used two input matrices composed of the “brain” data and the “cognitive/demographic” data detailed in Table 1A. Here, our “brain” data included the volume of each hippocampal subfield (matrix size 85×10). Our “cognitive/demographic” data contained age, sex, years of education, MMSE, APOE4 status and RBANS score for each subject (matrix size 85×6). Secondly, for the vertex-wise PLS, our “brain” data included either vertex-wise measurements of the SA or displacement in each hippocampus (matrix size 85×1215 or 85×1152). Our “cognitive/demographic” data contained age, sex, years of education, APOE4 status, and RBANS scores for each subject (matrix size 85×5).

Following a statistical protocol described in previous works (McIntosh and Lobaugh 2004; Krishnan et al. 2011; McIntosh and Mišić 2013; Zeighami et al. 2017; Patel et al. 2020), each LV was then tested statistically using permutation testing. First, row permutations (10,000 times) of the input “brain” matrix were subject to PLS in order to obtain a distribution of singular values with the hypothesis that a permuted “brain” matrix will eliminate the initial brain-cognitive relationships. From these permutations, a nonparametric p-value was calculated for each LV expressing the degree to which the singular value obtained from the original matrices was associated with chance. A p-value threshold of 0.05 was selected to demonstrate which LV had at least a 95% chance of not being associated with a random correlation in the original matrices.

Secondly, the degree to which each “brain” and “cognitive/demographic” variable contributes to a given LV was tested using bootstrap resampling. The rows of the “brain” and “cognitive/demographic” matrices were randomly sampled with replacement to generate 10,000 new sets of matrices, this time maintaining initial brain-cognitive relationships (in contrast to the permutation tests described above). Each bootstrapped pair was subject to PLS in order to create a distribution of singular vector weights for each variable. The ratio of the singular vector weight over the standard error of the weight, called a bootstrap ratio (BSR), represents the contribution and reliability of a “brain” variable. We used a BSR threshold of 2.58, analogous to a p-value of 0.01. Meanwhile, the distribution of singular vector weights of each cognitive variable is used to obtain a 95% confidence interval (Krishnan et al., 2011; McIntosh and Lobaugh, 2004; Nordin et al., 2018; Persson et al., 2014; Zeighami et al., 2017).

## 3. Results

### 3.1. Subfield-wise

Here, after testing linear, second- and third-order relationships between each subfield and age using the AIC, we observed a third-order relationship of volumes with age. Both hippocampi and all the subfields, except the right CA1, were significantly decreasing with greater age (Figure 2A). Most subfields expressed a slight and steady decrease until age 60, followed by an accelerated association until age 80. PLS analysis demonstrated that the left and right hippocampus and their subfields were each associated with one significant LV, explaining 93.7 % and 90.9 % of the covariance, respectively (Figure 2B). In the right hemisphere, decrease in all the subfield volumes was associated with older age (R = 0.38, 95% CI = [0.23,0.53]), female sex (R = 0.56, 95% CI = [0.45,0.67]), low education (R = -0.22, 95% CI = [-0.42,0.03]), low RBANS total score (R =-0.20, 95% CI = [-0.38,-0.02]) and low MMSE score (R =-0.24, 95% CI=[-0.41,-0.06]. Left LV1 identified a pattern in which decreased volumes in the CA1, CA4DG, subiculum, SRLM and ICV were associated with older age (R = 0.38, 95% CI = [0.23,0.55]), lower MMSE score (R=-0.18, 95% CI = [-0.36, 0]), and female sex (R = 0.49, 95% CI = [0.37,0.62]).

**Figure 2:**
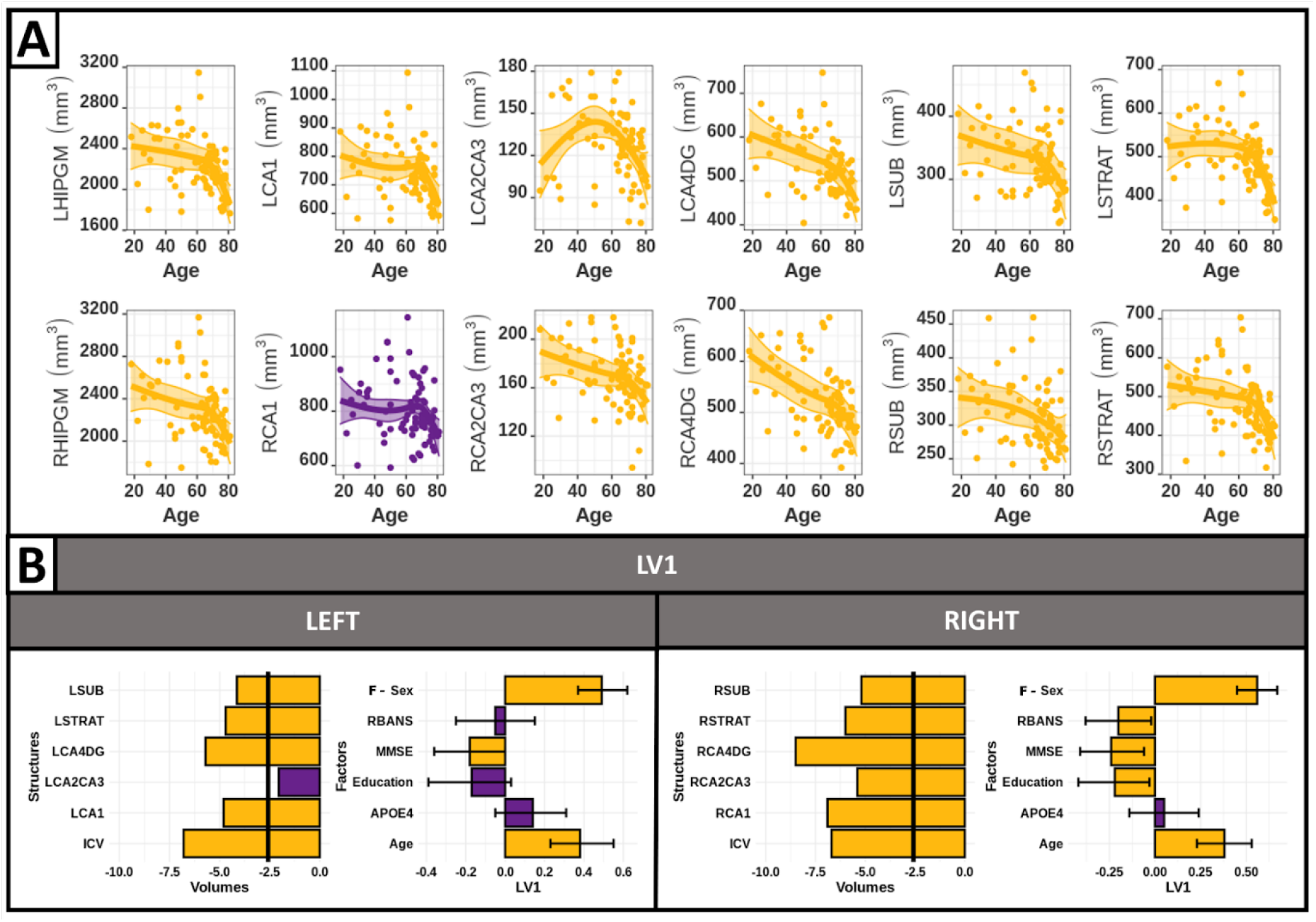
**A)** Third order relationships between age and the hippocampal subfields volumes. Purple: non significant; yellow: p<0.05 after Bonferroni correction. Supplementary table 2 illustrates t-values and uncorrected p-values. **B)** PLS between the left/right hippocampal subfield volumes and the demographic information identified one significant LV on each hemisphere (p<0.05). Right LV1 explained 93.7% and left LV1 explained 90.9% of the covariance. **Left side:** Bar plots describe the correlation of each subfield variable with the LV, yellow identifies variables significantly contributing to the LV. Black vertical line corresponds to a BSR threshold of 2.58. **Right side:** Bar plots describe the correlation of each demographic variable with the LV, with error bars denoting the 95% confidence interval, thus yellow identifies variables significantly contributing to the LV. Black vertical line corresponds to a BSR threshold of 2.58.

### 3.2. Vertex-wise

#### 3.2.1. Surface area

In Figure 3A, we use AIC at each vertex to show that, within the hippocampal structure, there are different types of age relationships. The right hippocampus had principally linear relationships with age in the body/tail, and a second or third-order relationship in the head of the hippocampus. The left hippocampus demonstrated a second-order relationship in the head, while the body and tail were best described linearly or with a third-order relationship with age. The vertices showing a significant effect of age are displayed in figure 3B. A significant linear decrease with age was found in the tail of the hippocampus, while the head of the hippocampus indicated significant second or third order relationships with age. Using these best models, we found that the maximum SA was largely found at 18 years old, except in the head of the hippocampus, which showed a maximum SA at ~60 years old (Figure 3C). Moreover, the minimum SA was reached at 81 years old, except in some local areas, like the uncus, which demonstrated a minimum SA at 18 years old. In Figure 3D, we plotted peak vertices of these different significant relationships with age. Of note, the second and third order curves corroborated that, for some vertices, the SA was preserved until the age of 60 years old.

**Figure 3:**
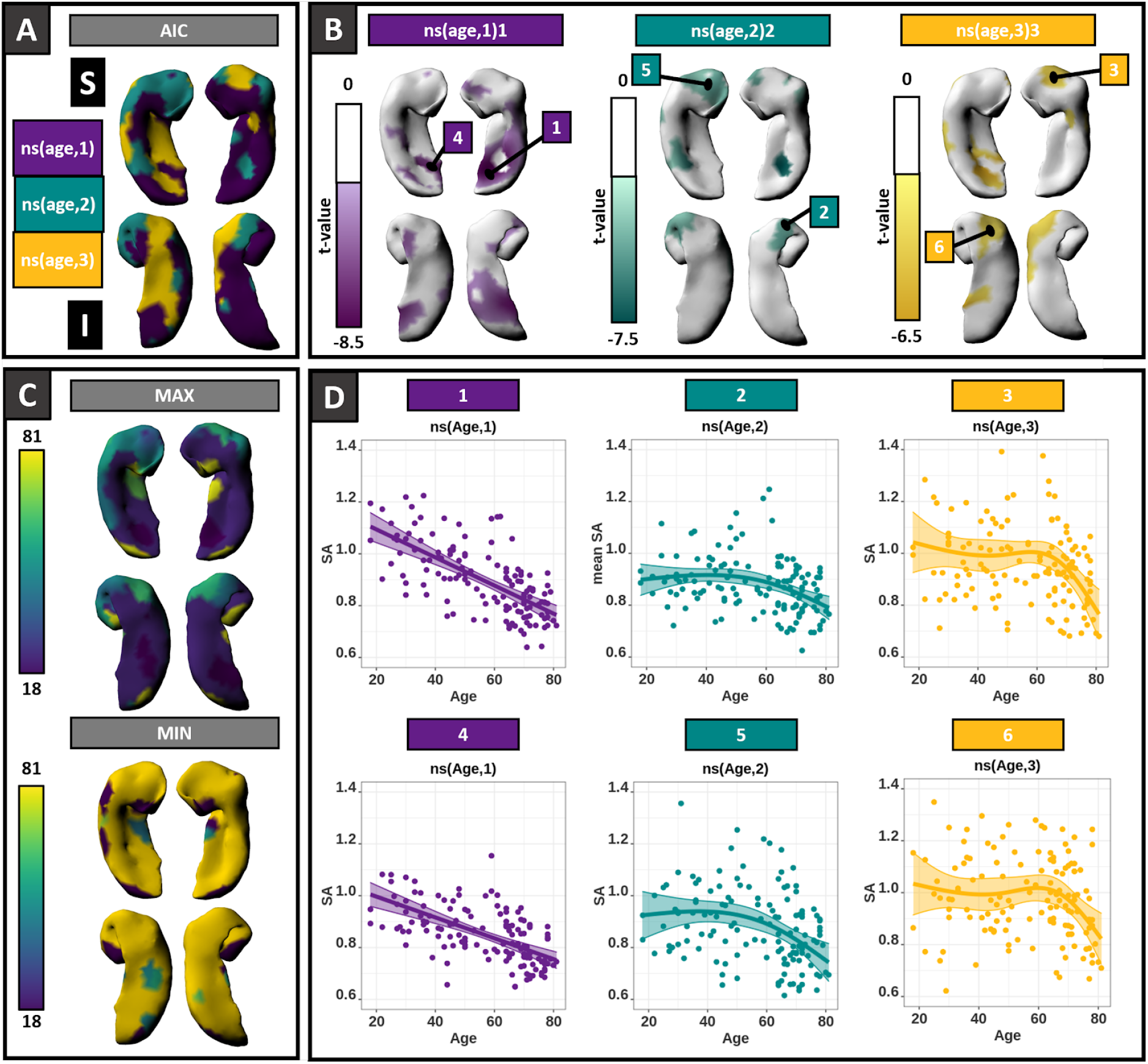
**A)** SA vertex-wise best fit age-relationship models as determined by AIC; purple for linear, cyan for second order and yellow for third order relationships with age; S: superior view; I: inferior view. **B)** Representation of significant age effects using the best model at each vertex (only higher order predictor shown) and the location of 6 significant peak vertices. t-value maps correspond to significant p-values (corrected at FDR 5%). **C)** Using the best model at each vertex, representation of the age for which SA was at its maximum or minimum. **D)** Plots of the SA of 6 peak vertices with age.

#### 3.2.2. Displacement

Here, we studied the relationship between age and displacement measures for each hippocampal vertex. In figure 4A, we used AIC at each vertex of the hippocampus, and found different types of age relationships. The models showing a significant effect of age are displayed in Figure 4B. A significant positive linear relationship with age was found in the inferior lateral hippocampus, while a significant negative relationship was found in the medial tail of the hippocampus. Significant positive second order best models were located in the lateral hippocampus, while negative second order models were situated in the right head and medial body of the hippocampus. Finally, a local positive third order displacement was contained in the left superior uncus, while a positive third order model was found in the left inferior uncus. Significant negative linear relationships were also observed, especially in the inferior medial tail of the hippocampus. Significant negative second order models were in the right superior head and along the medial body of the hippocampus. A third order negative displacement was located in the inferior left uncus. In general, the maximum displacement was found at an older age laterally and at a younger age in the superior body and medial inferior tail (see Figure 4C). Conversely, the minimum displacement was found in the superior tail and inferior head of the hippocampus. In Figure 4D, we plotted peak vertices of the different significant displacement change with age.

**Figure 4:**
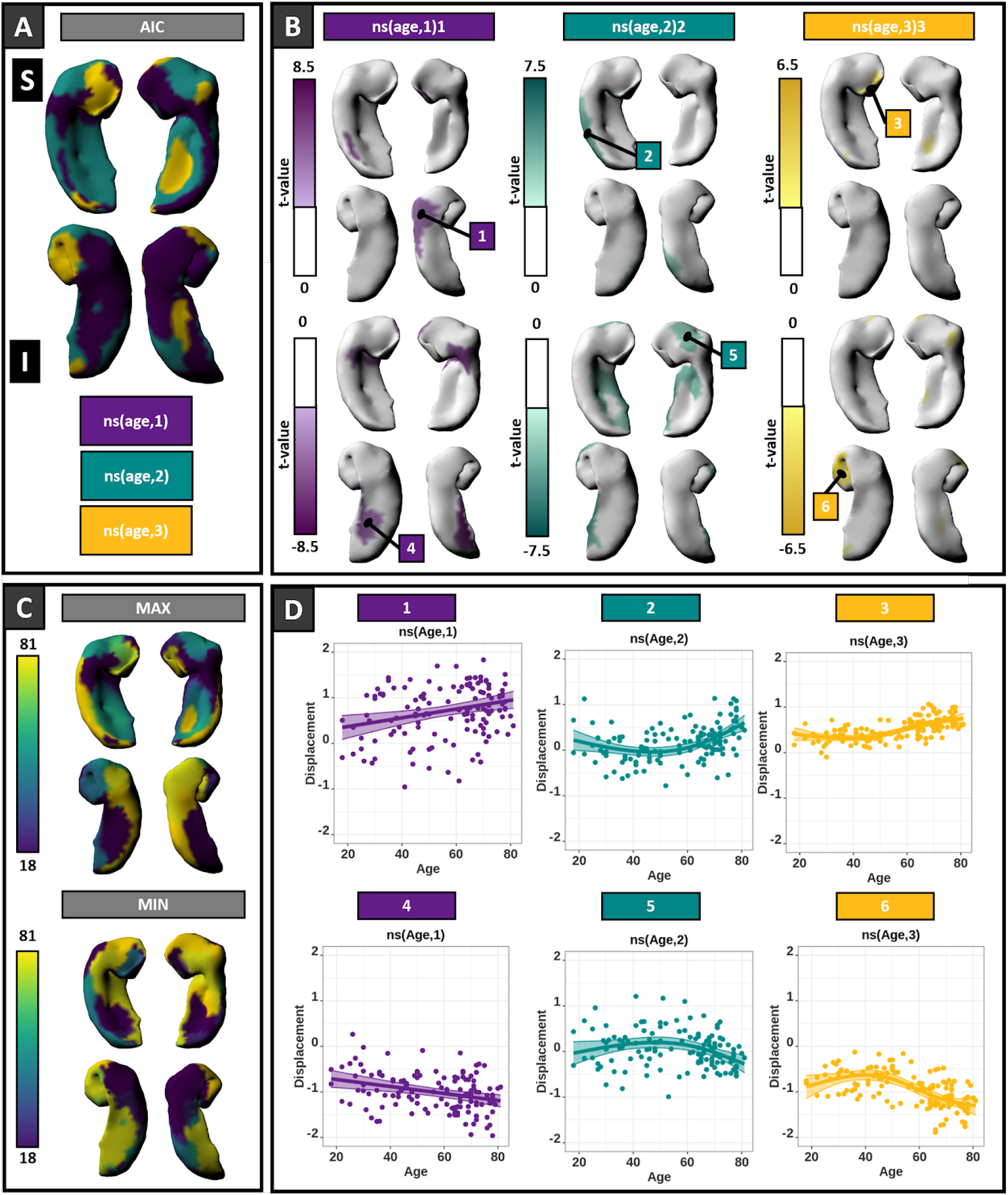
**A)** Displacement vertex-wise best age-relationship models from AIC; purple for linear, cyan for second order and yellow for third order relationships with age; S: superior view; I: inferior view. **B)** Representation of significant age effect using the best model at each vertex (only higher order predictor shown) and the location of 6 significant peak vertices. T-value maps correspond to significant p-values (corrected at FDR 5%). **C)** Using the best model at each vertex, representation of the age for which displacement was at its maximum or minimum. **D)** Plots of the displacement of 6 peak vertices with age.

### 3.3. Partial least squares analysis

#### 3.3.1. Surface area

For SA, PLS demonstrated three significant LVs, with the LV1 for the analyses for the right hippocampus explaining 79%, the LV2 for the analyses for the right hippocampus explaining 15% and the LV1 for the analyses for the left hippocampus explaining 68% of the covariance (Figure 5). Right LV1 represents a pattern in which decreased overall right SA is associated with older age (R = 0.39, 95% CI = [0.25,0.53]), female sex (R=0.55, 95% CI = [0.44,0.67]), low education (R=-0.22, 95% CI = [-0.40,-0.03]) and RBANS scores (R = -0.22, 95% CI = [-0.44,-0.04]). Left LV1 identifies a pattern in which decreased SA in the dorsal hippocampus is associated with older age (R = 0.41, 95% CI = [0.31,0.57]) and female sex (R=0 .33, 95% CI = [0.18,0.51]). Thus, similarly to the subfield-wise PLS, age and female sex are key factors explaining the hippocampal SA modifications. Also, the right hemisphere SA is more impacted by lower cognitive performance and levels of education. Significant effects of age impact regional SA, such that the right hemisphere with the right LV2 pattern shows an increased SA in medial tail and decrease SA in the uncus to be associated with younger age (R = -0.55, 95% CI = [-0.75,-0.20]).

**Figure 5:**
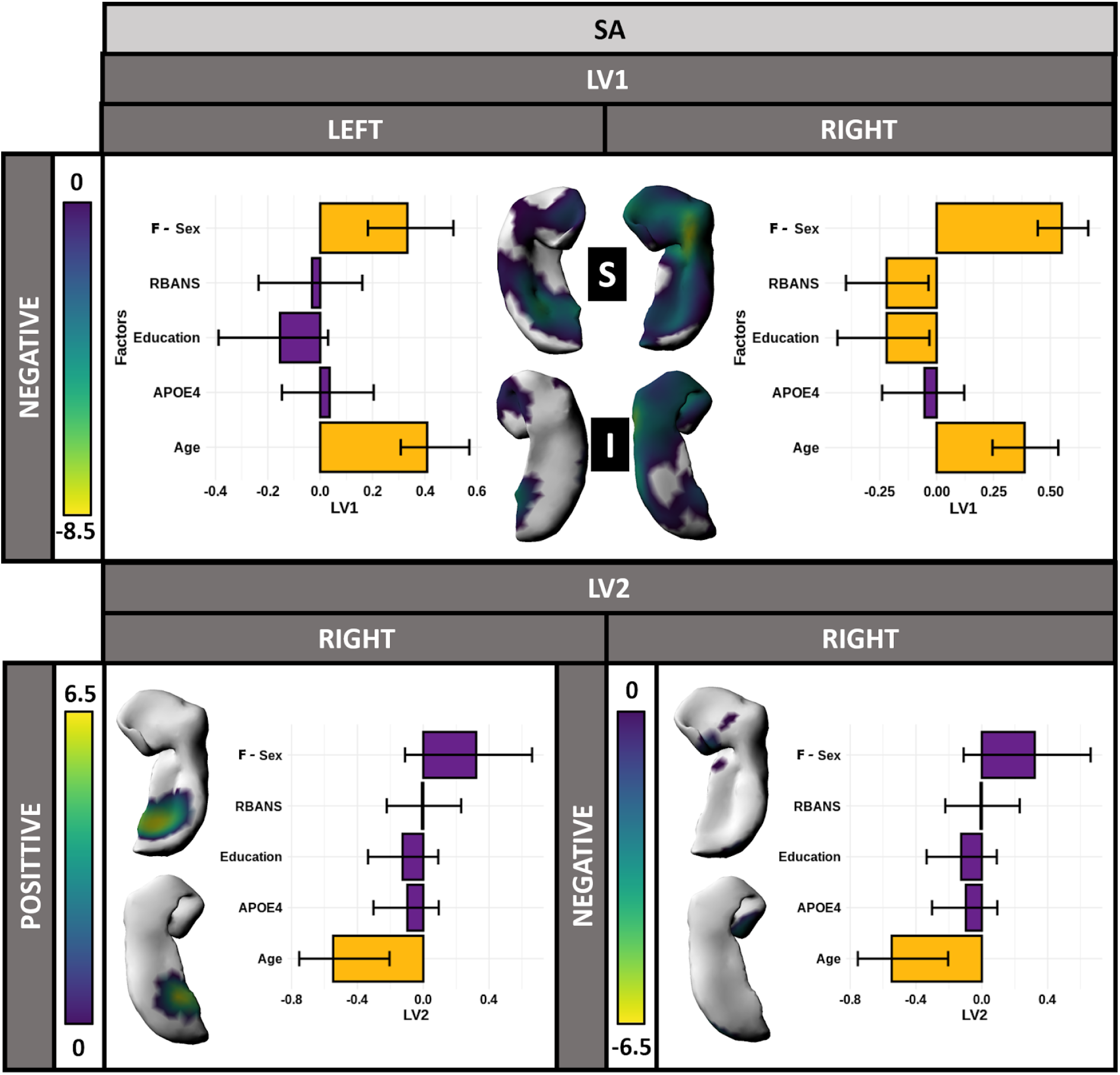
PLS analysis between the left/right SA and the demographic information identified two significant LVs on the right and one on the left (p<0.05); S: superior view; I: inferior view. Right LV1 explained 79%, right LV2 explained 15% and left LV1 explained 68% of the covariance. Bar plots describe the correlation of each demographic variable with the LV, with error bars denoting the 95% confidence interval; yellow identifies variables contributing significantly to the LV. For each LV, vertex wise BSR is plotted on the hippocampal surface and thresholded at 2.58 (p<0.01), describing the vertex wise SA patterns identified by each LV.

#### 3.3.2. Displacement

For displacement, PLS analysis demonstrated three significant LVs with the right LV1 explaining 52%, right LV2 explaining 29% and left LV1 explaining 77% of the covariance (Figure 6). Right LV1 showed a pattern in which outward displacement in the superior head and the medial tail, as well as inward displacement in the lateral body, was associated with younger age (R= -0.69, 95% CI = [-0.78,-0.63]). Left LV1 identified a pattern in which outward displacement in the superior body and inferior medial hippocampus, as well as inward displacement in the lateral hippocampus, was associated with younger age (R = -0.62, 95% CI = [-0.72,-0.54]), high RBANS score (R = 0.28, 95% CI = [0.13,0.44]) and higher levels of education (R= 0.23, 95% CI = [0.08,0.40]). In contrast to what was observed previously in the volumes and SA analyses, here, cognition and education appear to have an effect on the left hippocampus. Interestingly, a sex effect was only found in the right hemisphere with the right LV2, identifying a pattern in which inward displacement in the hippocampus head was associated with female sex (R = 0.41, 95% CI = [0.37,0.59]) and lower education levels (R = -0.24, 95% CI = [-0.43,-0.13]).

**Figure 6:**
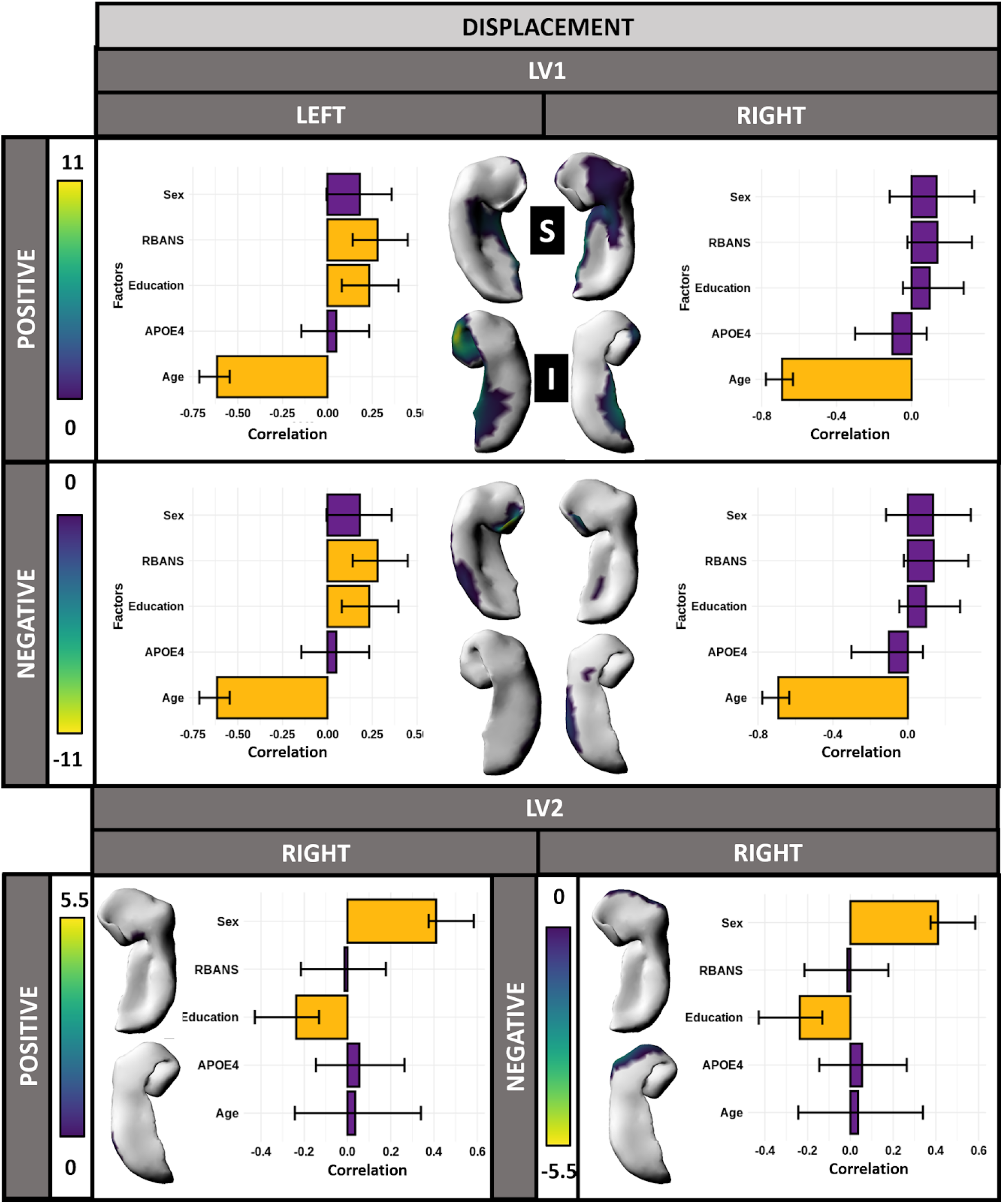
PLS analysis between the left/right displacement and the demographic information identified two significant LVs on the right and one on the left (p<0.05); S: superior view; I: inferior view. Right LV1 explained 52%, right LV2 explained 29%, and left LV 1 explained 77% of the covariance. Bar plots describe the correlation of each demographic variable with the LV, with error bars denoting the 95% confidence interval; yellow identifies variables contributing significantly to the LV. For each LV, vertex wise BSR is plotted on the hippocampal surface and thresholded at 2.58 (p<0.01), describing the vertex wise displacement patterns identified by each LV.

## 4. Discussion

The aim of this paper was to investigate the relationship between age and hippocampal structure throughout the healthy lifespan. To address this topic, we used standard subfield-wise volumetric as well as vertex-wise SA and displacement measurements. Subfield-wise investigation demonstrated third order relationships for all the hippocampal subfield volumes with age, highlighting an accelerated volumetric decrease after the age of 60. These results are in line with previous studies demonstrating a faster rate of hippocampal volumes decline during the sixth decade in healthy aging (Yang et al. 2013; Nobis et al. 2019; A. Bussy et al. 2020). Right CA1 volume appeared to be relatively less impacted by age, reproducing previous results from our group (Voineskos et al. 2015; Bussy et al. 2020; Amaral et al. 2018). Further, hippocampal volume analyses examining the influence of demographic factors emphasized that volume loss in most subfields (all except left CA2CA3) was strongly associated with older age and female sex, and to a lesser extent, in the right hemisphere, with lower cognitive performance and lower levels of education.

These results are in agreement with the literature; hippocampal subfield volumes are known to reduce during aging (Jack et al. 2015; de Flores, La Joie, and Chételat 2015).Moreover, lower levels of education (Noble et al. 2012) and low cognitive performance (O’Shea et al. 2016) are known to be risk factors for hippocampal atrophy. Also, we found that female sex was predominantly associated with low volumes and correlated with lower levels of education and cognition. Interestingly, hippocampal volumes have been found to be positively linked to associative memory in older women but not in men (Z. Zheng et al. 2017). Overall, this pattern of correlation between volumes and demographic characteristics was interesting but did not provide any information about the spatial specificity of these relationships since all subfields were implicated, except the left CA2CA3.

To investigate the spatial specificity of the impact of age in more depth, we ran vertex-wise analyses that demonstrated different patterns of age relationships throughout the hippocampal structure. The body/tail manifested a SA linear decrease, while the head suggested second and third order relationships with age. These findings illustrated a relative preservation of the anterior hippocampal SA with a peak during the sixth decade, while the posterior hippocampal SA gradually decreased across lifespan. Moreover, overall decrease of SA was linked to older age, female sex and, to a lesser extent in the right hemisphere, lower cognitive performance and lower levels of education. These results replicated the correlations found between the volumes and the cognitive/demographic factors described above. In addition, older age was specifically linked to lower SA in the right posterior hippocampus.

SA results are in agreement with previous findings demonstrating a stronger impact of aging in the posterior hippocampus (Malykhin et al. 2017; Langnes et al. 2020). Additionally, evidence of anterior/posterior (AP) anatomical and functional differences in the hippocampus have been highlighted by various studies. Namely, studies have indicated gradients of anatomical extrinsic connectivity with cortical and subcortical structures along the hippocampus long axis (Bryan A. Strange et al. 2014). Further, the posterior hippocampus appears to be specifically involved in spatial memory, while the anterior hippocampus is more involved in stress responses and emotional behavior (Fanselow and Dong 2010). Another recent study demonstrated that the anterior hippocampus showed higher functional connectivity with the anterior temporal lobe, orbitofrontal, inferior frontal gyrus, and premotor cortex, while the posterior hippocampus was more functionally connected to the medial and lateral frontal lobe, inferior parietal lobule, precuneus, and occipital lobes (Tang et al. 2020). This AP axis appeared to be related to hippocampal gene expression providing evidence of a molecular AP gradient (Vogel et al. 2020). Additionally, several previous and recent studies have suggested a hippocampal AP functional and anatomical gradient (B. A. Strange et al. 1999; Bryan A. Strange et al. 2014; Vos de Wael et al. 2018;Przeździk et al. 2019). Ultimately, our analyses support previous findings demonstrating a hippocampal AP axis. Also, our results indicate that the specific impact of age in the posterior hippocampus could illustrate that posterior hippocampal functions and connections might be more vulnerable than those in the anterior portion of the hippocampus.

In the present paper, analyses also indicated outward displacement with older age in the lateral hippocampus and inward displacement with older age in the medial hippocampus. Further, PLS analyses also strengthened this finding, demonstrating medial outward displacement and lateral inward displacement associated with younger age, and, to a lesser extent in the left hemisphere, higher levels of cognition and education. Interestingly, this pattern was distinct from the sex effect, which was found in the head of the hippocampus, in which inward displacement was associated with low education and female sex. Education has often been considered an influential factor in cognitive performance and hippocampal volume or microstructure (Piras et al. 2011) as well as a protective factor against mild cognitive impairment (Wada et al. 2018) and AD dementia (Meng and D’Arcy 2012; Groot et al. 2018). Gender inequality with regards to education is predictable since older women from our datasets were born in a socioeconomic period when they were less encouraged to pursue higher education than men (Permanyer and Boertien 2019). This could explain why female sex and low education are associated with age, and potentially contribute to our observed higher prevalence of women for hippocampal impairement-like measures. Furthermore, gender inequality regarding education could partly be responsible for the increased risk of developing AD in women (Beam et al. 2018).

Previously, medial/lateral (ML) gradient was found to correlate with an estimation of myelin content in the hippocampus, with the medial hippocampus having a higher myelin content (Vos de Wael et al. 2018). Further, a recent paper using a non-negative matrix factorization technique to cluster the hippocampus demonstrated that the medial hippocampus cluster was the most myelinated (Patel et al.2020). Similarly, intracortical myelin was found to be the highest in the subiculum, which is the most medial hippocampal subfield (DeKraker et al. 2018). Therefore, we hypothesize that the inward medial displacement with age observed in our study could potentially be associated with an age-related myelin decrease.

Interestingly, no effect of APOE4 genotype was established in our analysis, neither for the subfield volumes nor for the morphological measurements. Similarly, in a previous study performed in our laboratory, while APOE4 genotype demonstrated a trend for local inward displacement in the medial hippocampal head and outward displacement in the inferior head, these effects were not significant after 5% FDR correction (Voineskos et al. 2015). These outcomes are contradictory to previous findings showing increased atrophy rates in the hippocampus of APOE4 carriers (Shi et al. 2014; Li et al. 2016; Dong et al.2019). Nevertheless, other studies have found no evidence for a significantly greater rate of atrophy with increasing age (Taylor et al. 2014), nor even a stronger effect of age on hippocampal atrophy in the APOE4 non-carriers (Gonneaud et al. 2016; Aurélie Bussy et al. 2019).

Several previous studies that have parsed the hippocampus have used subjective and variable ways to define different regions of the hippocampus, both in terms of the manner by which the head, body and tail subregions were parsed (Duvernoy, Cattin, and Risold 2013; Malykhin et al. 2017) as well as the definitions of the anterior and posterior hippocampus. These methodological discrepancies could potentially explain some of the contradictory results, specifically that of anterior hippocampus volumes being more impacted by age, for example (Ta et al. 2012; Gordon et al. 2013). In the present paper, we believe that a methodological strength is that we performed our analyses with the goal of optimizing the statistics performed on our datasets; notably, the AIC method was used to select the best fit curves for each vertex (Akaike 1974; Mazerolle 2006). Accordingly, we found spatially well-defined and relatively bilateralized clusters of vertices, behaving either in a linear, second, or third order fashion, highlighting clear differences of age-related relationships throughout the hippocampus. Furthermore, in our PLS analyses, we demonstrated that the anterior hippocampal vertices already demonstrating decline in the first decades of life (expressing a linear decrease with age) were the ones highly associated with low cognition and education as well as with female sex. Moreover, SA relative preservation in the head of the hippocampus seems to be relatively well colocalized within the anterior CA1 subfield (see Figure 1), which might help to explain the relative preservation of CA1 volumes with age that we found when using subfield-wise measures. Nonetheless, this result seems to be in agreement with a previous study showing a similar finding on the CA1 preservation with age using a voxel-based morphometry technique (La Joie et al. 2010).

The entorhinal cortex (EC) is a structure known to connect a variety of cortical areas with the hippocampus. Sensory inputs are integrated in the superficial layers of the EC while layers II and III project to the hippocampus. Specifically, neurons from layer II project to the DG and CA3 through the perforant pathway, whereas layer III projects to the CA1 and subiculum through the mossy fibers (van Strien, Cappaert, and Witter 2009; Witter et al. 2017). The hippocampal circuitry ends with the subiculum projecting back to the layers V and VI of the EC, which has widespread cortical and subcortical projections. Selective vulnerability of neurons in layer II of the EC have been demonstrated in aging and AD (Gómez-Isla et al. 1996; Kordower et al. 2001; Stranahan and Mattson 2010). These results highlight a stronger age-susceptibility in layer II than in layer III. This could help explain our results since we have shown a higher age-related susceptibility in layer II projections, i.e. CA2CA3 and CA4DG subfields (Small et al. 2011), than in layer III projections, i.e. CA1 and subiculum. Indeed, some studies have demonstrated that regions with higher connection demonstrate selective vulnerability to disease-associated atrophy patterns (Seeley et al. 2009; Zhou et al. 2012; Shafiei et al. 2020).

Hippocampal development undergoes rapid growth and morphological modifications during the perinatal months. The hippocampal shape is likely influenced by the rate of neural progenitor migration in the developmental period and by neurogenesis in the DG until adulthood (Nowakowski and Rakic 1981;Kornack and Rakic 1999). Another study demonstrated that similar numbers of intermediate neural progenitors were found in the DG as well as comparable numbers of glia and mature granule neurons in cognitively normal individuals aged 14-79, indicating that neurogenesis persists throughout the course of healthy aging (Boldrini et al. 2018). Moreover, a study examining the impact of prematurity on the hippocampal shape demonstrated that at age 7, preterm children had greater hippocampal expansion in the AP direction and contraction along the medial border compared to full term children (Thompson et al.2014). Interestingly, this shows that perinatal development has the same effect of aging that we observed later in life (i.e an accentuation of the “C” shape). This leads us to hypothesize that the shape modification seen with age in our results is the continuation of the normal brain development and reorganization and not necessarily a marker of aging specific atrophy.

An important limitation to acknowledge is that this paper used cross-sectional data to investigate age-related hippocampal changes. While our results exhibit an interesting and promising understanding of hippocampal aging, we should keep in mind that longitudinal data would be necessary to validate these findings. Furthermore, the use of larger datasets could be relevant for such a validation study, as well as the use of higher resolution scans. Indeed, we previously demonstrated the impact of contrast and resolution on the hippocampal subfield volumes relationships with age (A. Bussy et al. 2020). Here, we decided to use T1w images because our morphometry technique is currently only applicable on T1w scans. However, further efforts to replicate these results using high resolution T2w images would be desirable.

To conclude, this paper aimed to describe the age-related relationships of the hippocampus volume and shape. Subfield-wise investigation demonstrated that the majority of the hippocampal subfields had accelerated volume decrease after 60 years old. Vertex-wise analysis provided more local information, showing a SA preservation in the anterior hippocampus, while the posterior hippocampus SA decreased linearly with age. Finally, displacement examination demonstrated a reinforcement of the hippocampus curvature with age.

## Acknowledgements

A. Bussy receives support from the Alzheimer Society of Canada. M. Chakravarty is funded by the Weston Brain Institute, the Canadian Institutes of Health Research, the Natural Sciences and Engineering Research Council of Canada and Fondation de Recherches Santé Québec.

## Supplementary

### Statistical models

Linear mixed-effects models (lmer from lmerTest_3.1-2 package in R 3.6.3) and natural splines (ns from splines package) were used to model linear, second, or third order age relationships for subfield-wise volumes. Akaike information criterion [AIC; (Akaike 1974)] was used to investigate the most appropriate age relationships across the multiple models. The model with the lowest AIC was selected and was considered to best fit the data (Mazerolle 2006). Sex, intracranial volume, APOE4 status, education, MMSE, and RBANS were included as fixed effects, while dataset was included as a random effect. *(1)*

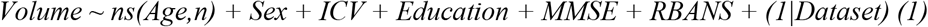

Linear mixed-effects models (vertexLmer from RMINC_1.5.2.2 package in R 3.6.3) and natural splines were used to model linear, second, or third order age relationships for vertex-wise measurements. AIC was used to investigate the most appropriate age relationships across the multiple models. The model with the lowest AIC was selected and was considered to best fit the data. Sex was included as fixed effects while dataset was included as a random effect. *(2)*

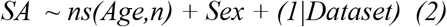

## Supplementary tables

**Table 1:** Initial demographics of the 178 participants included before motion QC.

|  | INITIAL |  |  |
| --- | --- | --- | --- |
|  | ADB<br>(n=66) | HA<br>(n=112) | Overall<br>(n=178) |
| **Age** |  |  |  |
| Mean (SD) | 69.8 (5.74) | 45.5 (16.4) | 54.6 (17.9) |
| Median [Min, Max] | 70.0 [56.0, 81.0] | 44.0 [18.0, 80.0] | 59.0 [18.0, 81.0] |
| **Sex** |  |  |  |
| F | 41 (62.1%) | 57 (50.9%) | 98 (55.1%) |
| M | 22 (37.9%) | 55 (49.1%) | 80 (44.9%) |

**Table 2:** T-values and p-values from linear mixed-effects models for the age relationship of the bilateral total hippocampus and each hippocampal subfield volume. Sex, intracranial volume, APOE4 status, education, MMSE, and RBANS scores were included as fixed effects, while dataset was included as a random effect but only the age values were reported. Significant p-values after Bonferroni correction were italicized and in bold.

|  | t-value<br>ns(Age,3)1 | t-value<br>ns(Age,3)2 | t-value<br>ns(Age,3)3 | p-value<br>ns(Age,3)1 | p-value<br>ns(Age,3)2 | p-value<br>ns(Age,3)3 |
| --- | --- | --- | --- | --- | --- | --- |
| **Left HIPGM** | -1.49 | -1.91 | -5.88 | 0.14 | 0.06 | ***1.35e<sup>-7</sup>*** |
| **Left CA1** | -0.28 | -1.72 | -4.03 | 0.78 | 0.09 | ***1.95e<sup>-4</sup>*** |
| **Left CA2CA3** | -0.28 | 1.20 | -3.54 | 0.78 | 0.23 | ***6.84e<sup>-4</sup>*** |
| **Left CA4DG** | -2.69 | -2.65 | -5.71 | 0.01 | 9.75e<sup>-3</sup> | ***2.64e<sup>-7</sup>*** |
| **Left subiculum** | -1.53 | -1.99 | -4.34 | 0.13 | 5.03e<sup>-2</sup> | ***4.55e<sup>-5</sup>*** |
| **Left SRLM** | -0.88 | -1.15 | -6.11 | 0.40 | 0.25 | ***3.34e<sup>-7</sup>*** |
| **Right HIPGM** | -1.62 | -2.08 | -4.53 | 0.12 | 0.04 | ***2.88e<sup>-5</sup>*** |
| **Right CA1** | 0.44 | -1.28 | -2.90 | 0.66 | 0.21 | 4.89e<sup>-3</sup> |
| **Right CA2CA3** | -1.84 | -1.93 | -3.70 | 0.08 | 0.06 | ***4.97e<sup>-4</sup>*** |
| **Right CA4DG** | -3.49 | -3.49 | -5.52 | ***9.66e<sup>-4</sup>*** | ***8.15e<sup>-4</sup>*** | ***5.20e<sup>-7</sup>*** |
| **Right subiculum** | -2.03 | -1.17 | -3.04 | 0.05 | 0.25 | ***3.55e<sup>-3</sup>*** |
| **Right SRLM** | -1.21 | -1.60 | -4.75 | 0.24 | 0.11 | ***1.95e<sup>-5</sup>*** |

